# Warped Bayesian Linear Regression for Normative Modelling of Big Data

**DOI:** 10.1101/2021.04.05.438429

**Authors:** Charlotte J. Fraza, Richard Dinga, Christian F. Beckmann, Andre F. Marquand

## Abstract

Normative modelling is becoming more popular in neuroimaging due to its ability to make predictions of deviation from a normal trajectory at the level of individual participants. It allows the user to model the distribution of several neuroimaging modalities, giving an estimation for the mean and centiles of variation. With the increase in the availability of big data in neuroimaging, there is a need to scale normative modelling to big data sets. However, the scaling of normative models has come with several challenges.

So far, most normative modelling approaches used Gaussian process regression, and although suitable for smaller datasets (up to a few thousand participants) it does not scale well to the large cohorts currently available and being acquired. Furthermore, most neuroimaging modelling methods that are available assume the predictive distribution to be Gaussian in shape. However, deviations from Gaussianity can be frequently found, which may lead to incorrect inferences, particularly in the outer centiles of the distribution. In normative modelling, we use the centiles to give an estimation of the deviation of a particular participant from the ‘normal’ trend. Therefore, especially in normative modelling, the correct estimation of the outer centiles is of utmost importance, which is also where data are sparsest.

Here, we present a novel framework based on Bayesian Linear Regression with likelihood warping that allows us to address these problems, that is, to scale normative modelling elegantly to big data cohorts and to correctly model non-Gaussian predictive distributions. In addition, this method provides also likelihood-based statistics, which are useful for model selection.

To evaluate this framework, we use a range of neuroimaging-derived measures from the UK Biobank study, including image-derived phenotypes (IDPs) and whole-brain voxel-wise measures derived from diffusion tensor imaging. We show good computational scaling and improved accuracy of the warped BLR for certain IDPs and voxels if there was a deviation from normality of these parameters in their residuals.

The present results indicate the advantage of a warped BLR in terms of; computational scalability and the flexibility to incorporate non-linearity and non-Gaussianity of the data, giving a wider range of neuroimaging datasets that can be correctly modelled.

## 1. Introduction

Big data has become more widely available in neuroimaging (UK Biobank, ENIGMA, ABCD study, PNC, among others) [1], [2], [3], [4]. This has ignited a renewed interest in modelling normal brain development, to estimate quantitive brain-behaviour mappings and capture deviations from such models to derive neurobiological markers of different psychiatric disorders. These neurobiological markers could move us closer towards individualized and precision medicine [5]. Until now, the neurobiological markers for psychiatric disorders have been mostly developed with studies that used classifiers trained in a case-control setting. Counter-intuitively, an increase in sample size has shown to reduce the accuracy of classifying cases from controls for psychiatric disorders [6]. One of the main reasons for this decrease in accuracy has been posed to be the heterogeneity in the participants both biologically and behaviorally, which can only fully be captured by a large data set [6]. Normative modelling is an emerging method used to understand this heterogeneity in the population. Similar to growth charts in pediatric medicine, which describe the distribution of height or weight of children according to their age and sex, normative models can be used to model the distribution of neuroimaging derived phenotypes in a population, including the mean and centiles of variation [7], according to age, gender, or other demographic or clinical variables [8]. The deviations from this normative range can be quantified statistically, for example as Z-scores, which have been linked to several psychiatric disorders [7], [9], [10], [11], [12], [13].

Although promising, there are still certain challenges in performing normative modelling on big neuroimaging data. First of all, Normative models have been mainly developed using Gaussian process regression. [14]. Gaussian process regression is flexible and accurate, but a drawback is its computational complexity, which is governed by the need to compute the exact inverse of the covariance matrix. This inversion scales poorly with an increase in data points [15]. Therefore, using these models on large datasets requires extensive computational power and is often not feasible (typically beyond a few thousand subjects). Furthermore, most normative models assume the modelled distribution is Gaussian. However, distributions diverging from Gaussianity are frequently found in specific neuroimaging modalities. These non-Gaussian signals cannot be accounted for using a standard normative model based on Gaussian process regression. We argue that modelling non-Gaussianity is important in general and is frequently overlooked by the neuroimaging community in that most regression methods used in practice –often implicitly– assume Gaussian residuals. Thus, there is a need to develop methods that can flexibly handle the computational demand and non-Gaussianity of big data sets.

In this paper, we propose a next-generation framework based on Bayesian linear regression (BLR) to address these challenges. We introduce an extension of the BLR method for accurately modelling non-Gaussian distributions using a likelihood warping technique, giving a warped BLR model. The new framework has several benefits over previously used methods: (i) A BLR model can use a linear combination of non-linear basis functions (such as B-splines) which can be considered to provide a low-rank approximation of the Gaussian process regression models [16]. However, the BLR model has considerably better computational scaling, since the complexity of the model is fixed according to a set of basis-functions. Therefore, the model can be scaled much more easily to large datasets. Furthermore, a set of model coefficients can be estimated that can easily be shared without the need to share the data (e.g. to compute a cross-covariance matrix for new data points), thus making it easier to make predictions on new datasets. (ii) The non-Gaussianity of the residuals can be modelled by the flexible warping of the Gaussian function, which gives more options to model different types of neuroimaging data that cannot be accurately modelled using a standard BLR. (iii) Efficient model selection criteria are naturally defined for the warped BLR through the marginal likelihood and can be calculated in closed form. The marginal likelihood gives a balance between model complexity and model fit. This can aid in choosing the optimal model for a specified imaging modality.

We will demonstrate this model by testing it on different types of neuroimaging data derived from the UK Biobank dataset. The UK Biobank dataset has several magnetic resonance imaging (MRI) imaging modalities, including structural and functional brain data. With over 40,000 participants’ MRI data from 40 to 80 years old, this provides a rich set of different neuroimaging data and defines a benchmark for future population-based studies. In this work, we will present the warping function and recommend how to use it for several data modalities. First, we give an illustrative example using image-derived phenotypes (IDPs), which are convenient and widely used summary measures of brain function and structure [17]. Specifically, we will show a detailed example of estimating a normative model for white matter hyperintensities (WMHs). WMHs have been shown before to demonstrate quite non-Gaussian behaviour [18], and are therefore a good example where the warped BLR could be preferred over the B-spline BLR. Second, we show the scalability of this method by performing a whole-brain analysis for certain diffusion tensor imaging (DTI) measures. We use DTI measurements, as there are large associations with age and we expect certain non-linear and non-Gaussian trends in the data [19].

Finally, we want to evaluate the link between brain imaging abnormality scores and behaviour. Therefore, deviations from normal brain functioning are associated with cognitive functioning. The deviations are captured by Z-scores, which are shown to correlate with measures of intelligence in the UK Biobank dataset, such as; numerical memory, reaction time and visual memory.

In summary, the main contributions of the paper are to give: (i) a new comprehensive framework for big data normative modelling; (ii) the introduction of the novel methodological approach for modelling non-Gaussian response variables; (iii) an extensive and didactic evaluation of this framework on the UK Biobank cohort and (iv) a demonstration of the ‘Predictive Clinical Neuroscience software toolkit’ (PCNtoolkit) for big data normative modelling. Ultimately, we hope this paper will give deeper insight into the application of normative models on different types of neuroimaging modalities.

## 2. Materials and methods

### 2.1. Sample

All the data used came from the UK Biobank imaging dataset [1]. Full details on the design of the study and the preprocessing steps can be found in subsequent papers [17], [20]. Briefly, the data used contains around 10,000 participants of the 2017 release and additional longitudinal data of around 5,000 subjects of the 2020 release. The participants were between 40 and 80 years of age, with around 47 % males.

In this study, two types of analyses were performed using different datasets. For the first analysis, a dataset containing IDPs was used. For consistency with existing work, the IDPs were processed using FUNPACK [21], which is an automatic normalisation, parsing and cleaning kit, developed at the Wellcome Centre for Integrative Neuroimaging. The IDPs include three main imaging modalities: structural, functional and diffusion brain imaging. Among these IDPs, there are very gross measures, such as the total amount of brain volume, to more detailed measurements, such as the connectivity between two brain regions. In total 819 neuroimaging IDPs were used for subsequent analysis, see B.1 for the list of IDPs used. Furthermore, we also tested our model on another set of IDPs; 150 FreeSurfer measures, which were preprocessed with FreeSurfer v6.1.0, using a *T* 2-weighted image where available, see B.1 for the list of the FreeSurfer measures used.

For the second analysis, a whole-brain model was built, using voxel-wise fractional anisotropy (FA) and mean diffusivity (MD) measures. The data were processed using the UKB pipelines; including the DTI fitting tool DTI-FIT and a tract-based spatial statistics (TBSS) style analysis, which gave us the skeletonised DTI files. In total, around 10,000 participants with dMRI-scans passed the quality control applied by the UK Biobank [17]. Afterwards, we added two extra exclusion criteria. First, participants were removed if their Z-score of the discrepancy between the dMRI image and the structural T1 image was higher than three, based on data-field 25731 in the UK Biobank. Second, participants were removed if their Z-score of the number of outlier slices was higher than three, which is a reflection of the movement of the participant during the scan, based on data-filed 25746-2.0 in the UK Biobank. For the covariates we used age, gender and dummy coded site variables.

### 2.2. Cognitive data

We used the cognitive phenotypes that were extracted from the UK biobank using FUNPACK [21] to evaluate the cognitive associations with the deviations from the normative model. These measures are derived from the 13 cognitive tests present in the UK Biobank, see the UKB showcase. The tests were administered using a touchscreen questionnaire and included numerical memory, reaction time, fluid intelligence, visual memory and prospective memory. Later other tests that measured executive function, declarative memory and non-verbal reasoning were added [22]. For full details on the different cognitive tests applied in UK Biobank see [23]. An overview of all the measures used in this study is presented in the supplementary E.6.

### 2.3. Normative model formulation

We use a flexible normative modelling framework to model different types of neuroimaging data. We have *N* subjects with brain data 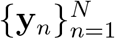, each of dimension *D* (e.g. the number of voxels or IDPs) and acquired from one of *S* different scanning sites. We use **Y** to denote an *N* × *D* matrix containing these variables, where *y*_*nd*_ denotes the *n*-th subject and *d*-th neuroimaging variable. Since the neuroimaging variables are estimated separately here, we simplify the notation by using **y** to denote the vector of observations from a single variable and *y*_*n*_ for a single observation. In general, we want to predict the distribution of the value for each voxel or brain region, the dependent variable (**y**), from a set of covariates 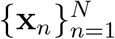 (e.g. age, gender or site), the independent variables. In this paper, we adopt a straightforward approach to model nonlinear relationships, by applying a basis expansion to the independent variables. A common approach is to use polynomials, but these can be a poor choice, as they can induce global curvature [24]. Here we apply a common B-spline basis expansion (specifically, cubic splines with 5 evenly spaced knot points), although other approaches are also possible. We denote this expansion by *ϕ*(**x**), with **Φ** an *N* × *K* matrix containing the basis expansion for all subjects. In the applied model, *y* is assumed to be the result of a linear combination of the B-spline basis function transformation plus a noise term:

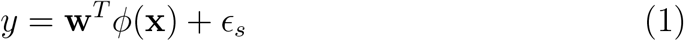

With **w** the estimated vector of weights and 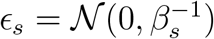 a Gaussian noise distribution for site s, with mean zero and a noise precision term *β*_*s*_ (i.e. the inverse variance). All the noise precision terms from the different sites will be combined in a vector ***β*** and the site precision matrix **Λ**_***β***_, which has ***β*** along the leading diagonal and is the inverse of the site covariance matrix **Λ**_***β***_ = **Σ**_***β***_^*−*1^. Note that we allow the noise precision to vary across sites in order to accommodate inter-site variation along with site-specific intercepts (i.e. dummy coded site regressors in the design matrix). We have shown previously that this approach provides an efficient way to accommodate site effects in normative modelling [25].

Following similar derivations as given by Huertas et al. [16], we consider a BLR model, placing a Gaussian prior over our model parameters *p*(**w**|***α***) = 𝒩 (**w**|0, **Λ**_***α***_^*−*1^), with ***α*** the hyper-parameters that the weights depend on. The Gaussian prior is assumed to have a mean zero and a precision matrix **Λ**_***α***_, with the precision matrix the inverse of the covariance matrix **Σ**_***α***_ = **Λ**_***α***_^*−*1^. As shown in Huertas et al. [16], **Λ**_***α***_ can be quite general, but here we use a simple isotropic precision matrix **Λ**_***α***_ = *α***I**. The Gaussian prior choice allows us to compute the posterior distribution of **w** in a closed form:

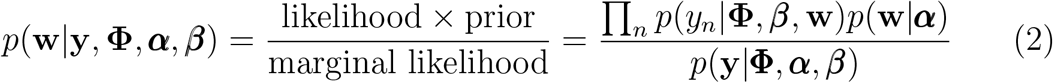

The posterior for each subject can then be found using the standard derivations of the posterior [26]:

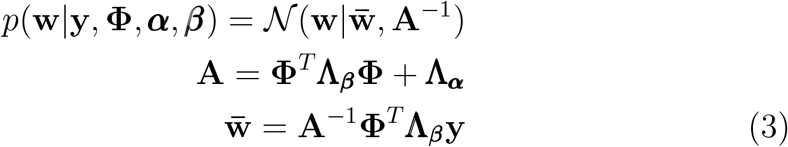

We use a Type II maximum likelihood approach (i.e. empirical Bayes), optimizing the denominator of the posterior to find the optimal hyper-parameters ***α*** and ***β***. This gives an automatic trade-off between model fit and model complexity. The marginal likelihood is maximized by minimizing the negative log likelihood (NLL):

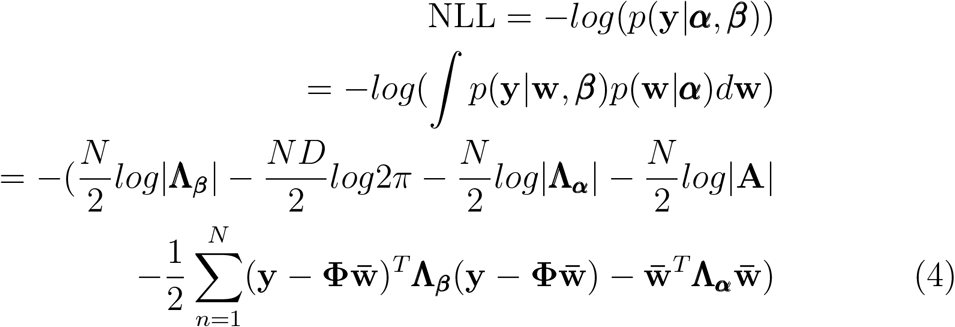

The optimal hyper-parameters ***α*** and ***β*** are often estimated using a conjugate gradient optimisation of the NLL, where the derivatives can be computed directly. However, here we used Powell’s method to fit the hyper-parameters. Powell’s method is a derivative-free method, which in this case is faster, because computing the derivatives of the marginal likelihood with respect to the hyper-parameters is computationally very expensive. Finally, the predictive distribution is given by:

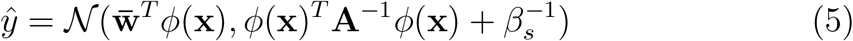

#### 2.3.1. Likelihood warping

In order to model non-Gaussian error distributions, we employed a ‘warped’ likelihood [27]. This involves applying a non-linear monotonic warping function *ϕ*_*i*_ to the input data during the model fit, with the index *i* indicating a different warping function (e.g. SinArcsinh, Box-Cox etc.). This is similar to the classical statistical technique of variable transformation, but has the advantage that the parameters of the transformation are optimised during model fitting, to provide the optimal mapping that ensures that model residuals have a Gaussian form. The warped functions are chosen such that they have a closed form inverse and are differentiable, which has several benefits: first, non-Gaussian data can be mapped (i.e. warped) exactly to better match Gaussian modelling assumptions or the predictions can be warped back to the original non-Gaussian space; second, it allows inference, prediction and computation of error measures all in closed form; finally, we can construct compositions of functions from the invertible monotonic warping functions that can greatly improve the expressivity of the model in transforming non-Gaussian distributed data **y** to a Gaussian form, **z**, where inference is straightforward [28]. This is done by applying a compositional warping function *ϕ* to the observations **y**:

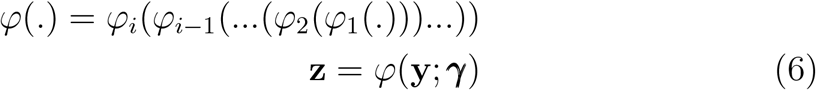

With ***γ*** denoting the hyper-parameter(s) of different warping functions. The warping transformation allows us to compute error measures in the warped space and to describe the deviations of subjects under a Gaussian error distribution in the form of pseudo Z statistics, even if the original data distribution is non-Gaussian.

The optimal hyper-parameters (***α, β*** and ***γ***) are calculated by minimizing the warped NLL. The warped NLL can be found by accounting for the change of variables in the probability density function [28]:

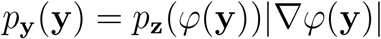

With ∇*φ*(.) the Jacobian of the transformation, which is diagonal and therefore we can simplify as a product of the individual terms:

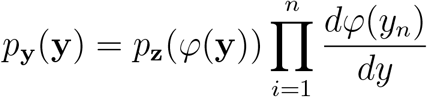

If we take the negative log of this equation the warped NLL will remain the same as equation 4, except for replacing the **y** by the transformed *ϕ*(**y**) and the inclusion of the Jacobian term that takes the change of volume induced by the warping into account, thereby ensuring a valid probability measure, for details see [28]:

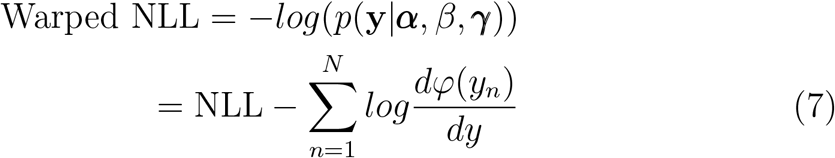

#### 2.3.2. Computational complexity

The optimization of the hyper-parameters is controlled by the minimization of the warped NLL. The warped NLL consists of the basic BLR NLL term and the log-derivatives of the warping *ϕ*_*i*_ functions, which are known in closed-form by construction. The complexity of the warped BLR model is then roughly the same as the classic BLR. However, the warped NLL is optimized for an extra hyper-parameter ***γ***, which could lead to the presence of more local minima, making the optimization process slightly slower [28].

#### 2.3.3. Warped composition function

Different elementary functions can be used to create the warped composition function *ϕ*. For this paper, we test affine, Box-Cox and SinhArcsinh transformations and compositions of these transformations:

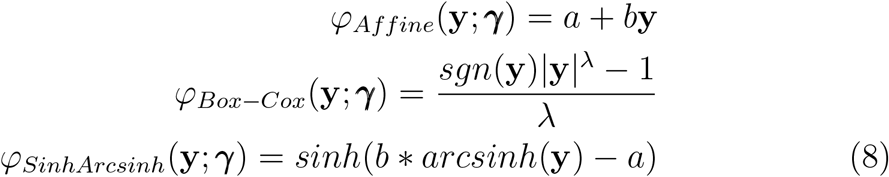

With ***γ*** the respective parameters of the different warping functions. For the SinArcsinh warping we also applied a reparametrization [29], as this empirically gave more stable results:

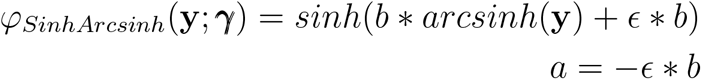

### 2.4. Model selection

We evaluate the models using different model selection criteria. First, we calculate the explained variance (EV) of the model. It is expected that the gain in fit for the warped BLR will be highly dependent on the flexibility of the model. Therefore, the Bayesian Information Criterion (BIC) is also considered:

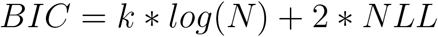

Which penalises for model complexity. Here *N* denotes the number of participants in the training set, NLL the negative log-likelihood. *k* is the number of free parameters. Note that we use the marginalized from of the NLL, which already takes into account the number of estimated coefficients. Therefore, the BIC only needs to be corrected for the added complexity of the degrees of freedom of the model (i.e. the parameters that are not integrated out). For the standard BLR this is two, one for the precision over the weights and one for the precision over the noise (*α* and *β* respectively). For the warped SinArcsinh BLR two extra degrees of freedom are added for the shape parameters (*a* and *b*). The BIC gives a good trade-off between the extra flexibility found in the warped BLR model and the better fit of the model. Finally, the mean standardized log-likelihood (MSLL) is used as a third model criterion. The MSLL takes into account the mean error and the estimated prediction variance.

### 2.5. Deviance scores and correlation to cognitive phenotypes

We want to find a statistical estimate of how much each participant deviates from the normal range. This is done by computing a Z-score for each subject *n*, also denoting explicitly the dependence on each voxel or IDP *d*:

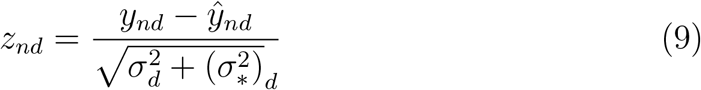

With, ŷ_*nd*_ the predicted mean and *y*_*nd*_ the true response. Normalized by 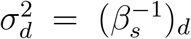 the estimated noise variance (i.e. reflecting variation in the data) and 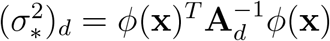 the variance attributable to modelling uncertainty for the *d*-th voxel. For the warped statistic, we compute the Z-scores in the warped (i.e. Gaussian) space. The true response variables are warped to the Gaussian space to ensure the underlying assumption of normality is satisfied by the construction of the warping functions.

Afterwards, to ensure our model can also be applied for behavioural and clinical estimations, we look at the correlations between the Z-scores from the IDPs and the whole brain analysis, and the cognitive scores of the UK Biobank. For the IDPs, we directly correlate the Z-scores and the cognitive phenotypes through a Spearman correlation. For the whole-brain analysis, we first make a summary statistic of the Z-scores by calculating the extreme value distribution. We model the extreme value distribution by looking at the mean of the top 1% of the deviations across the whole brain [10]. The extreme value statistics give the largest deviations per subject from the normal pattern, which have shown to be strongly correlated to behaviour [10], [30]. Afterwards, we apply a principal component analysis (PCA) on the cognitive phenotypes to give a one-factor solution. This first component has been shown to be correlated to the ‘general’ factor of cognitive ability or the ‘g-factor’ [31]. Lastly, we compute the Spearman coefficient between the first principal component and the summary deviation score.

## 3. Results

### 3.1. Performance of the warped Bayesian linear regression model for IDPs

All the statistical analyses were performed in Python version 3.8, using the PCNtoolkit. The BLR algorithm from the PCNtoolkit was chosen for all experiments. We considered age, binary gender and binary site ID within the covariance matrix. We used a standard BLR or we transformed the age covariate with a B-spline of order three with three knots. The Powell method was selected for the optimizer. We randomly split the dataset into 50% training and 50% test and reported all the error metrics on the test set. In the PCNtoolbox, several warpings can be chosen depending on the imaging modality one wants to model. We tested several warping functions (affine, Box-Cox and SinhArcsinh) and compositions of these warping functions. Preliminary testing showed that the SinhArcsinh warping gave the best fit compared to the alternatives evaluated. Therefore, in this paper, only the results of the SinhArcsinh warping are presented.

In figure 1, Bland-Altman plots are shown comparing the standard BLR and the B-spline BLR. The figure presents different model selection criteria: MSLL and BIC (EV can be seen in supplement figure A.8). The plots demonstrate that for most IDPs a non-linear B-spline BLR model performs better than a standard BLR. Indicating that non-linearity is a key component that should be accounted for in modelling neuroimaging data.

**Figure 1:**
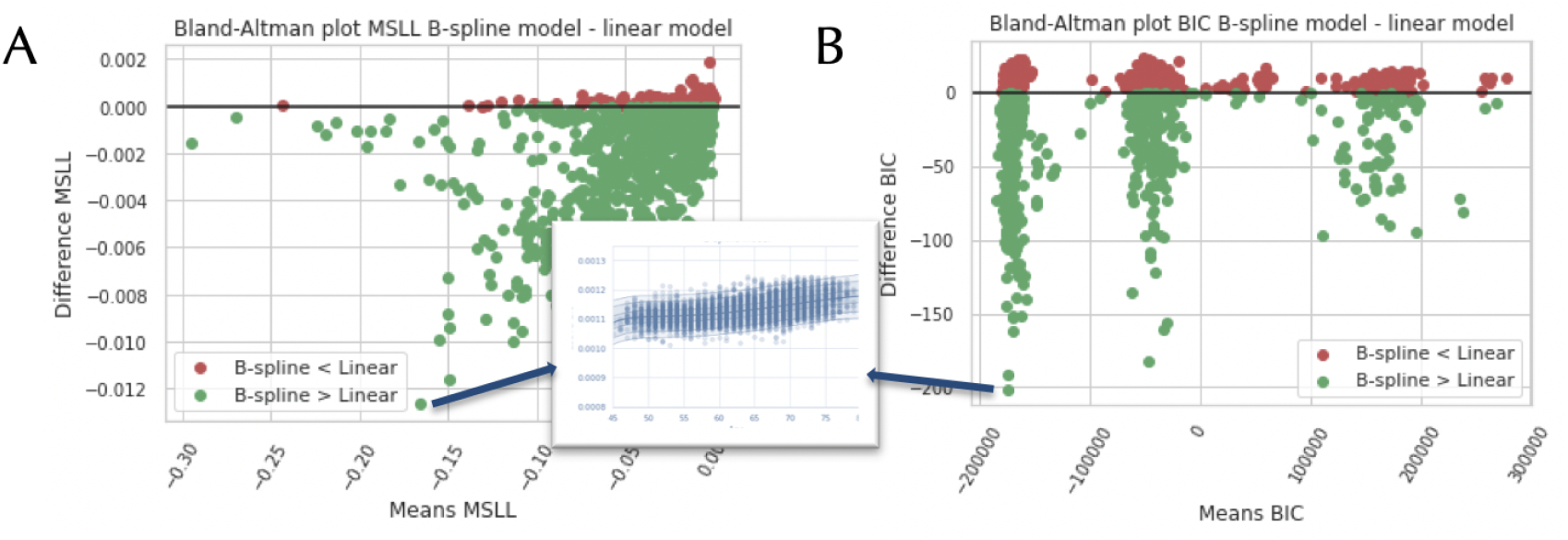
Bland-Altman plots comparing the standard and B-spline Bayesian Linear Regression (BLR) models, using Image-Derived Phenotypes (IDPs). Each dot indicates one IDP. The models are compared according to the following model selection criteria: the Mean Standardized Log Loss (MSLL) (A) and the Bayesian Information Criteria (BIC) (B). The green colour indicates a better fit for the non-linear B-spline model compared to the linear model. We also plotted a zoomed-in view of the model fit for one of the IDPs.

In figure 2, Bland-Altman plots are shown that compare the B-spline BLR and the warped BLR models for all IDPs, using the MSLL and BIC (EV can be seen in supplement figure A.8). We also plotted the difference in absolute values of the skewness and kurtosis. In figure 3, the same plots are shown for the FreeSurfer measures. We included them separately, as they were preprocessed separately (i.e. we did not use the IDPs provided by UK Biobank and instead ran the Freesurfer reconstructions manually). The plots show that for specific IDPs the warped BLR performs better than the B-spline BLR. When we examined these IDPs more closely, it was noted that they demonstrated distinct non-Gaussian behaviour. An example of such behaviour is given down below with the WMHs (white matter hyperintensities). In the supplementary table C.3, we provide a summary of some of the results for different IDPs that can help inform which neuroimaging modalities are best modelled with the warped BLR. For an indication of the effect sizes of the model selection criteria for the different model settings, see supplementary tables D.4 and D.5. Note also that the MSLL and EV do not clearly reflect differences in the shape of the predictive distribution. For example, for the IDPs, there is no average difference between the warped and non-warped model (Fig. 2 panel A and supp. fig. A.8 panel B), yet the warped model consistently yields a predictive distribution –and resultant Z-score distribution– that is less (or equivalently) skewed and kurtotic (Fig. 2 panels C and D).

**Figure 2:**
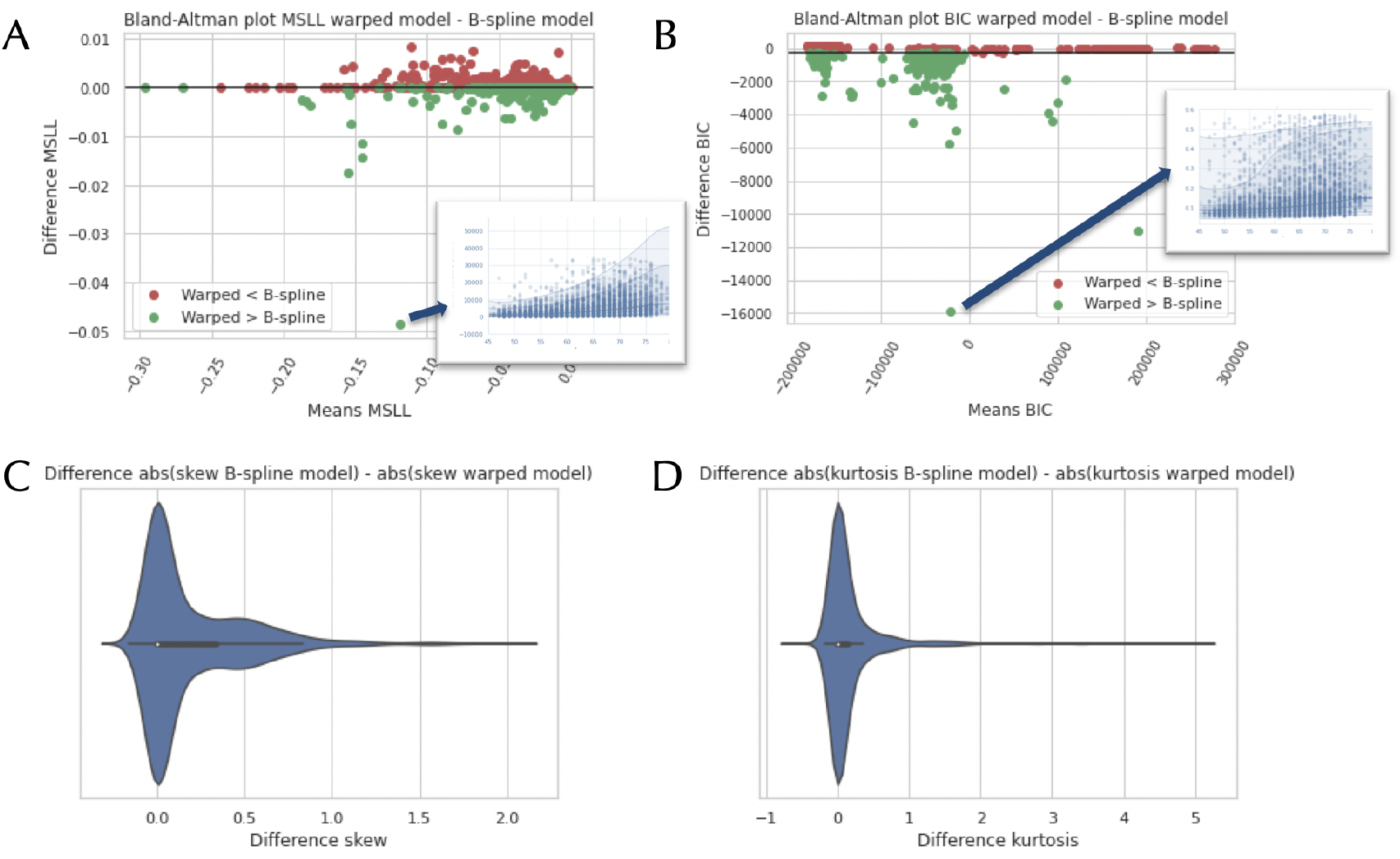
Bland-Altman plots comparing the B-spline and warped Bayesian Linear Regression (BLR) models, using Image-Derived Phenotypes (IDPs). The models are compared according to the following model selection criteria: the Mean Standardized Log Loss (MSLL) (A) and the Bayesian Information Criteria (BIC) (B). The green colour indicates a better fit for the warped model compared to the B-spline model. We also plotted a zoomed-in view of the model fit for two of the IDPs. On images C and D, we show the difference in absolute values of the skewness and kurtosis between the B-spline and warped model. A more positive value indicates that the B-spline model had a higher skewness or kurtosis than the warped model.

**Figure 3:**
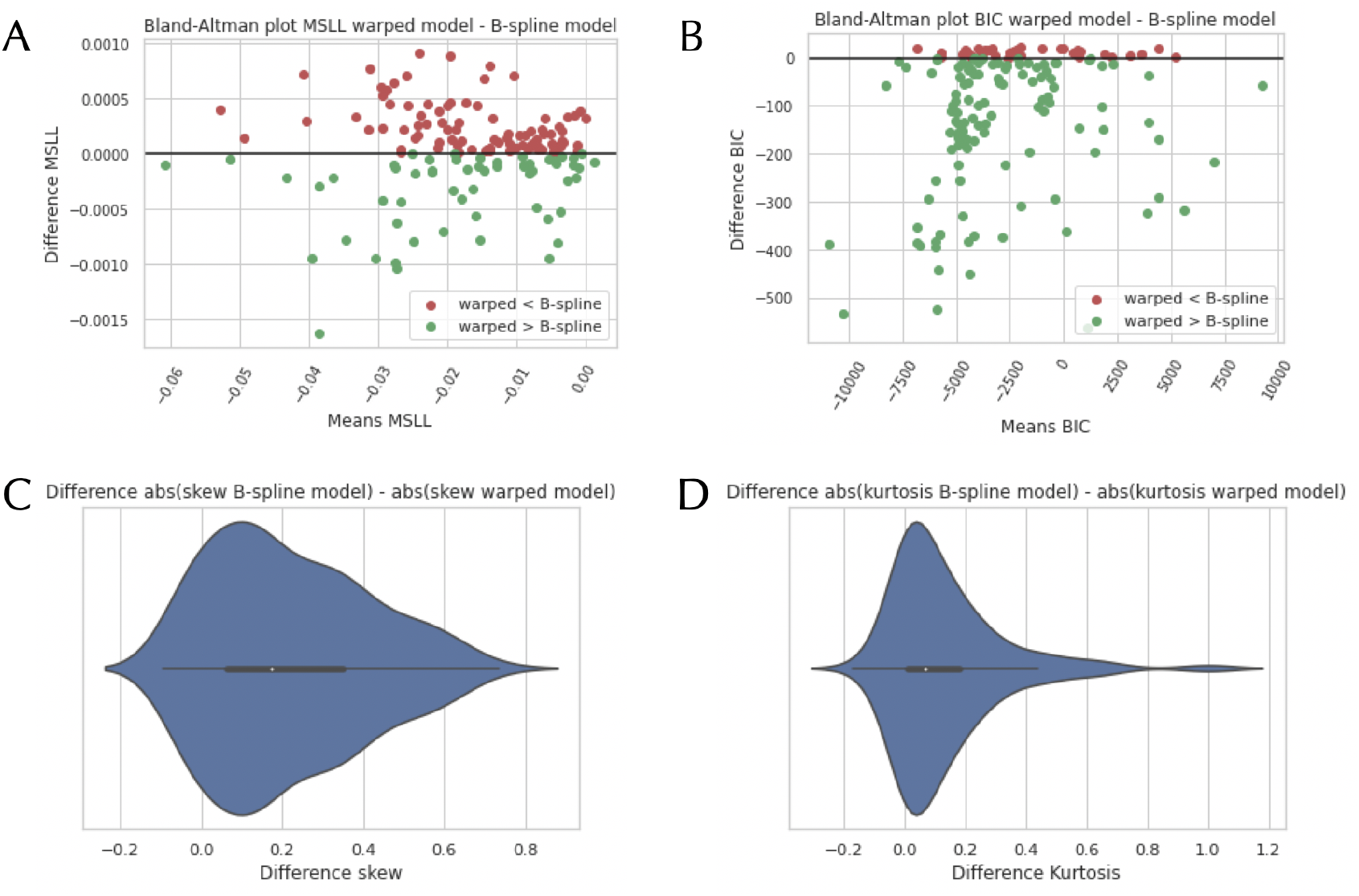
Bland-Altman plots comparing the B-spline and warped Bayesian Linear Regression (BLR) models, using the FreeSurfer measurements. The models are compared according to the following model selection criteria: the Mean Standardized Log Loss (MSLL) (A) and the Bayesian Information Criteria (BIC) (B). We also plotted a zoomed-in view of the model fit for one of the IDPs. On images C and D, we show the difference in absolute values of the skewness and kurtosis between the B-spline and warped model. A more positive number means a better fit for the warped model compared to the B-spline model.

In figure 4 and 5, we show the results of an illustrative analysis predicting WMH load across ageing to demonstrate how the performance of the warped BLR model compares to a B-spline BLR. The figures show the B-spline BLR and warped BLR results for WMHs at one-time point and the longitudinal data of two-time points. The results demonstrate that (i) the non-linearity of the data is sufficiently captured with a B-spline transformed BLR (ii) the WMHs show a distinctly non-Gaussian variance pattern, which is better predicted by the warped BLR. Thus, indicating that if the data has a non-Gaussian distribution for the residuals a warped BLR is preferred over a B-spline BLR.

**Figure 4:**
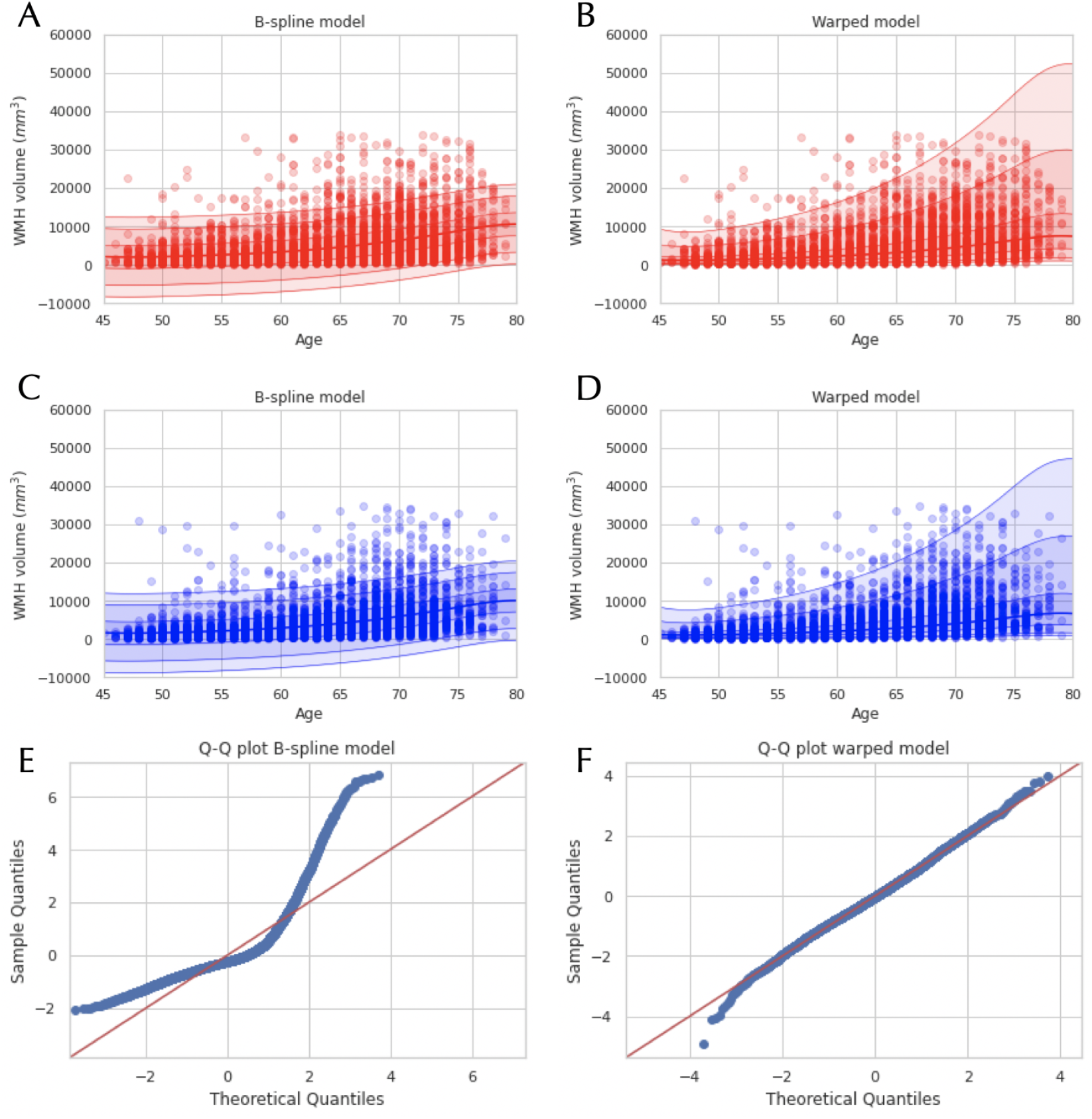
White matter hyperintensities (WMHs) modelled as a function of age using a Bayesian Linear Regression (BLR) model. Images A and C demonstrate the model fit using a regular Gaussian B-spline BLR, for the female and male cohorts respectively, both visualizing the mean prediction and the centiles of variation for the WMHs. Images B and D show comparable fits for a SinArcsinh warped BLR, for the female and male cohorts respectively. In images E and F quantile-quantile (QQ) plots of the two models are shown, demonstrating a better fit for the data using a warped BLR model.

**Figure 5:**
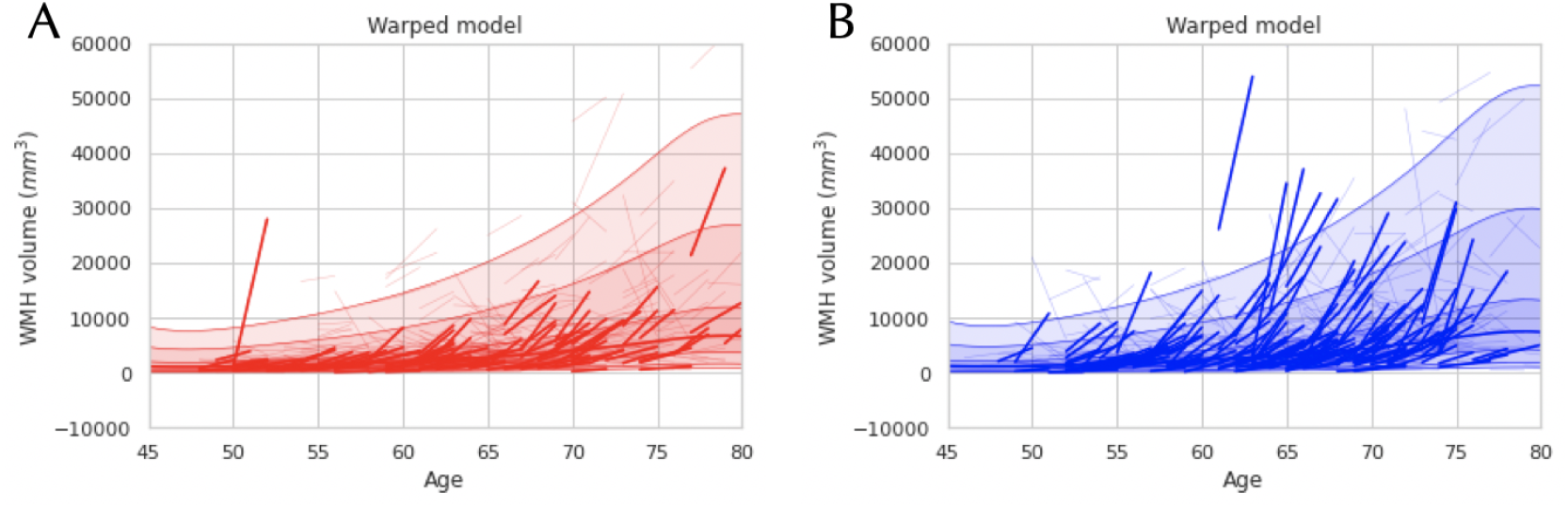
Here the longitudinal follow-up data of the WMHs is plotted for females (A) and males (B), using a SinhArcsinh warped BLR model.

#### 3.1.1. Correlation deviance scores WMHs and cognitive phenotypes

We also wanted to correlate the warped BLR model output of the WMHs to behavioural variables to ensure that the model can be used for behavioural predictions. We loaded all cognitive phenotypes available in UK Biobank according to the FUNPACK categorization, including: reaction time, numeric memory, prospective memory etc. (for a full list of the cognitive phenotypes used, see the supplementary table E.6). We calculated the deviance Z-scores according to formula 9. Afterwards, we calculated the Spearman correlation between the cognitive phenotypes and the Z-scores. Numeric memory (ID: 4259, ‘Digits entered correctly’) was modestly but significantly correlated with the warped Z-scores: *ρ* = −0.0331, *p* = 0.0262. In other words, if a participant’s WMH deviation from normal development increases the number of correctly remembered digits drops.

Lastly, to illustrate the value of normative models in a longitudinal context, we tested for an association between change in WMHs and change in cognitive phenotypes of the longitudinal data to see if WMH load is correlated to cognitive decline. We performed a statistical Wilcoxon rank-sum test on the participants’ cognitive phenotypes contrasting subjects that have a difference in the Z-scores *>* 0.5, which corresponds to a difference in half a standard deviation, versus the participants that do not. Intuitively, this contrasts individuals who are following an expected trajectory of ageing with those who deviate from such a trajectory. Highly significant associations were found with the reaction time (ID: 404, ‘Duration to first press of snap-button in each round’) *W* = 5.5641, *p* < 0.001 and with the Trail Making Test (ID: 6771, ‘Errors before selecting correct item in alphanumeric path (trail #2)’) *W* = 8.3105, *p* < 0.001. The results show an association between the change in cognition and the change in WMH deviance scores.

### 3.2. Scalability to a whole brain voxelwise based analysis

For the follow-up analysis, we evaluated the warped BLR approach on a whole-brain level for two DTI imaging modalities (FA and MD). The results of these two modalities were very similar and therefore we will only present the results for FA here. We separated the entire dataset into 80% training data and 20% testing data. First, we computed the time complexity per model fit (e.g. for one voxel) with varying number of subjects using the B-spline BLR model setting and compared it to the Gaussian process regression setting (Figure 6). This demonstrates the clear computational advantage of the BLR setting for the whole brain analysis.

**Figure 6:**
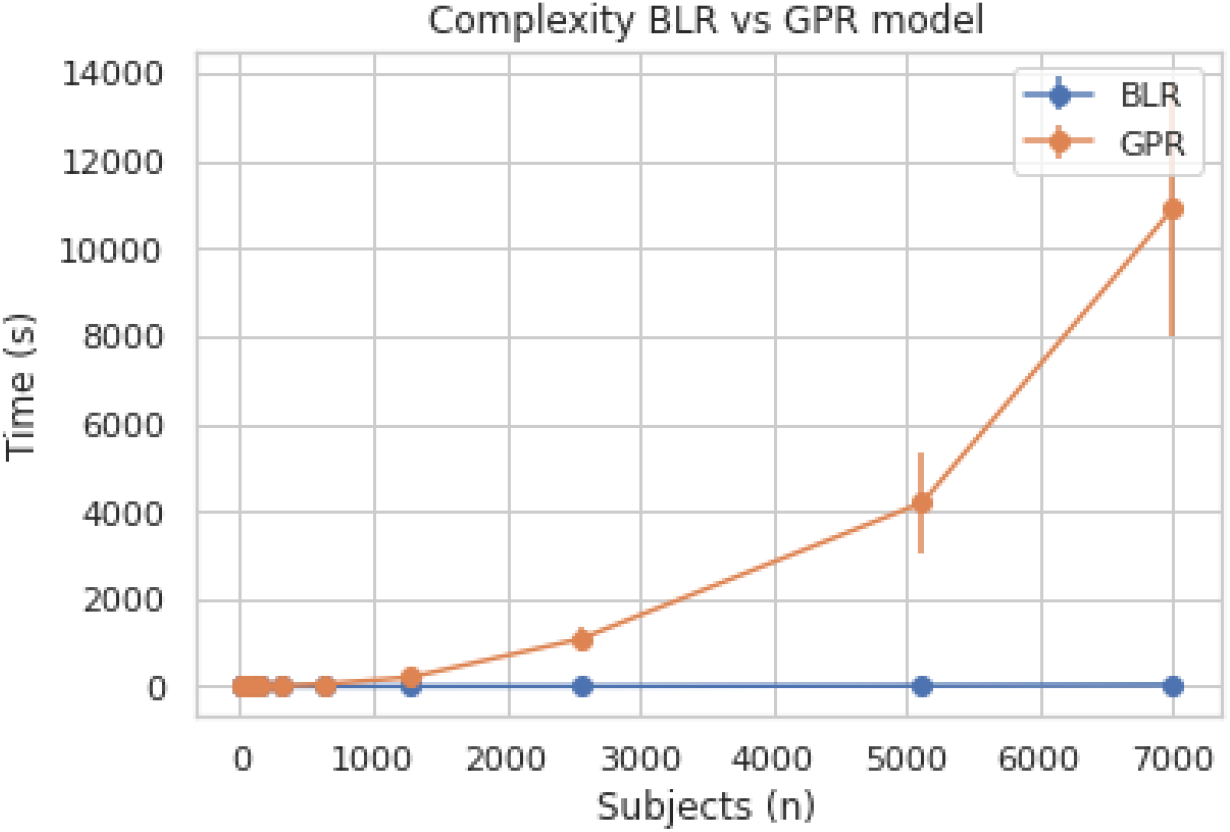
Computational complexity comparison between the Bayesian linear regression (BLR) model setting and the Gaussian process regression (GPR) model setting, giving the mean and the standard error (SE) over ten runs.

Afterwards, we tested different model settings for the imaging modalities including a standard BLR, B-spline BLR and a SinhArcsinh warped BLR. Figure 7 shows the comparative results in a Bland-Altman plot for the FA dataset (which were similar for the MD dataset). The figure presents the EV, MSLL and the BIC for the B-spline BLR and the warped BLR. These results are consistent with the IDPs in that according to the EV and MSLL, the models perform quite similarly for most voxels. Although, we would argue that these measures are not necessarily sensitive for the added benefit of the warping of the likelihood, which will mostly affect the predictions in the outer centiles. For the BIC the results demonstrate that the warped BLR is preferred for certain voxels. The voxels where a warped model is favoured generally showed more non-Gaussian behaviour.

**Figure 7:**
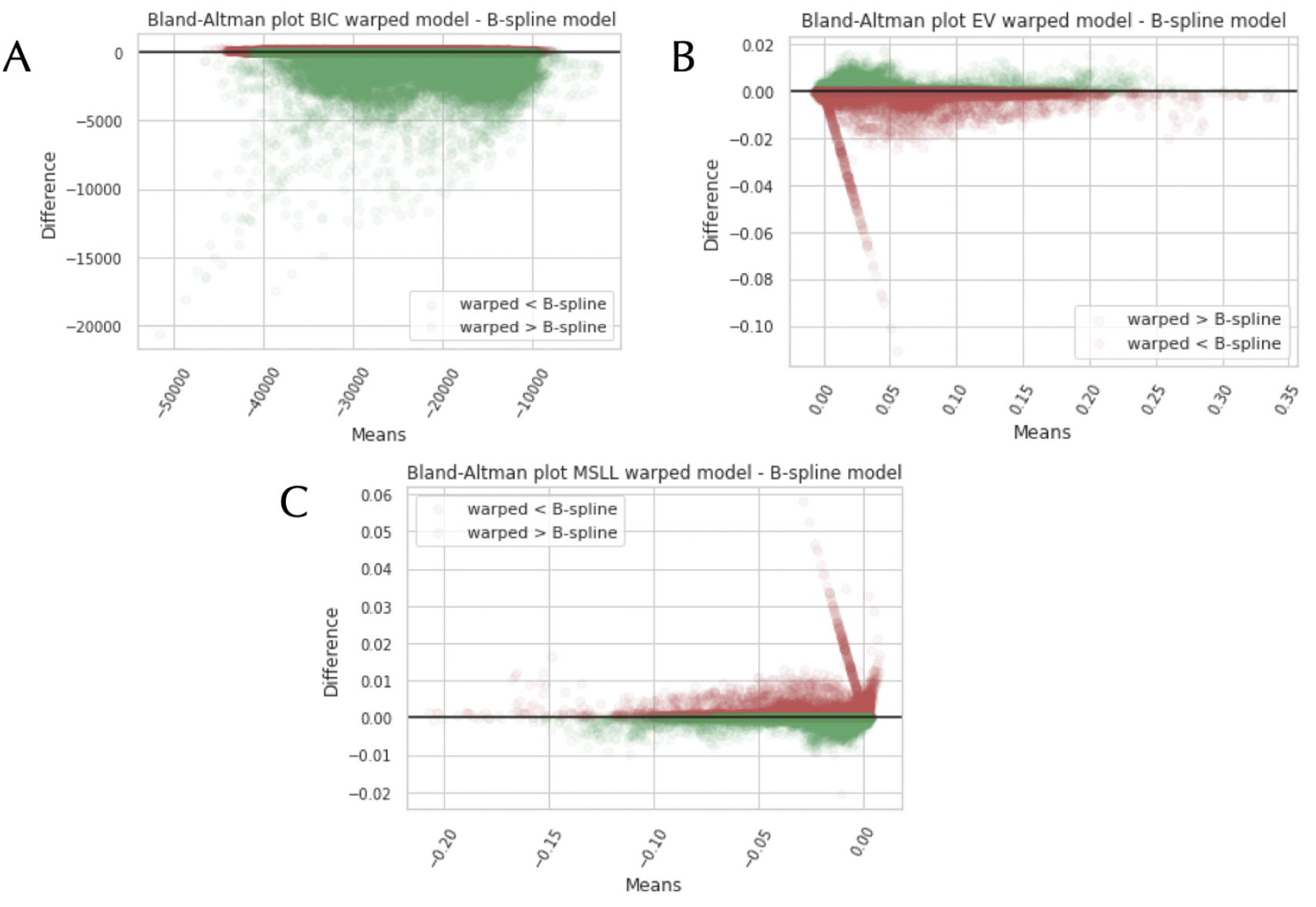
Bland-Altman plots comparing the warped Bayesian Linear Regression (BLR) model to the B-spline BLR model, using Fractional Anisotropy (FA) data. The comparison is done according to the following model selection criteria: The Bayesian Information Criteria (BIC) (A), the Explained Variance (EV) (B), and the Mean Standardized Log Loss (MSLL) (C). The green colour indicates a better fit for the warped BLR.

**Figure A.8:**
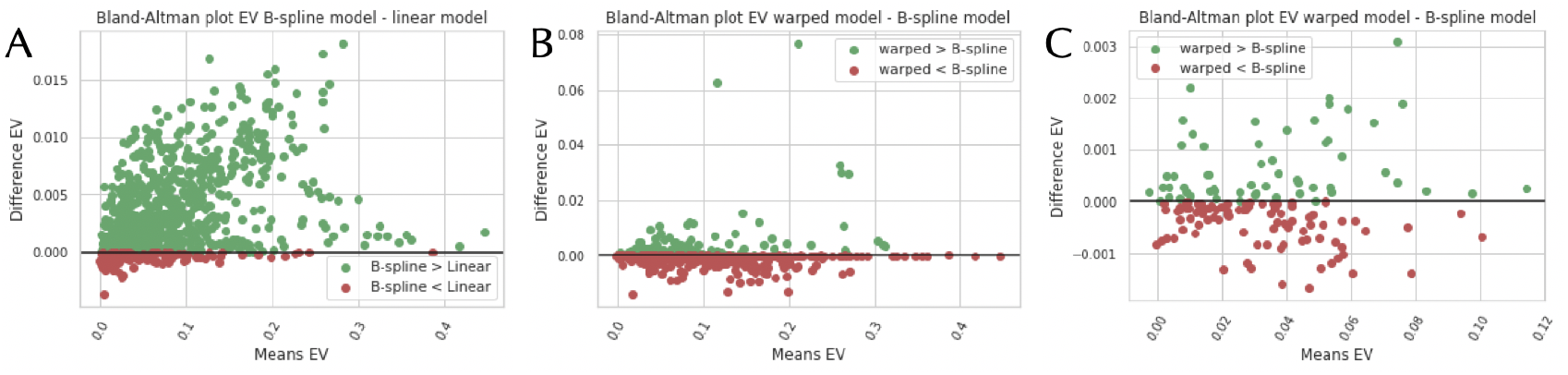
Bland-Altman plots of the Explained Variance (EV): Figure A shows the comparison of the linear and B-spline model, using the IDPs. Figure B shows the comparison of the warped and B-spline model, using the IDPs. Figure C shows the comparison of the warped and B-spline model, using the FreeSurfer measurements.

Finally, We used a paired-sample t-test, pairing the whole brain results (EV, MSLL and BIC) of the different models to estimate the difference between performance measures of the warped and non-warped BLR. For MD the following effect sizes were found: *EV* : *d* = 0.33, *MSLL* : *d* = 0.003 and *BIC* : *d* = −0.79. For FA the following effect sizes were found: *EV* : *d* = 0.028, *MSLL* : *d* = 0.017 and *BIC* : *d* = 0.55. We can see that the difference between the methods is small. Indicating that the B-spline BLR and the warped BLR model are quite similar in their model fit for MD and FA.

#### 3.2.1. Correlation deviance scores DTI and cognitive phenotypes

Finally, we correlated the Z-scores of the whole brain warped BLR model for the MD dataset to the cognitive phenotypes. First, we scaled the cognitive data and performed a principal component analysis. We selected the first component, which explained 29% of the variance in the data. Afterwards, we made a summary score of the Z-scores for each participant by looking at the largest deviations, which in the limit should follow an extreme value distribution [32]. We fitted a generalized extreme value distribution to the top 1% of the absolute Z-scores of each subject. Subsequently, we computed a Spearman correlation between the extreme values and the first principal component of the cognitive phenotypes, which gave *ρ* = 0.158, *p <* 0.001. The results demonstrate a clear correlation between the warped deviations from normal development and the cognitive phenotypes. This relationship will be explored further in future studies.

## 4. Discussion

In this paper, we presented a next-generation framework to scale normative models for large population-sized datasets based on warped Bayesian linear regression (BLR). Normative models can capture the heterogeneity in the population and model individual deviations from normal brain development. We demonstrated that the shift in normative modelling to a B-spline BLR with a likelihood warping gives several benefits. In this study we showed that: (i) Compared to Gaussian process regression, it is computationally much less demanding and is therefore scalable to big datasets. (ii) The non-linearity of the model, incorporated by the B-spline, improves the fit and out of sample predictions for most variables. (iii) Non-Gaussianity of the data can be naturally included due to the incorporation of the likelihood warping in the algorithm, which allows for a wider range of datasets to be accurately modelled. (iv) Model selection criteria based on the marginal likelihood, such as the BIC, can be calculated in closed form and therefore a trade-off between model fit and model complexity can be chosen optimally from the training data, without cross-validation. (v) The deviations scores from normal brain development can be meaningfully related to behaviour. Furthermore, we demonstrated the use of the normative model with the warped BLR on different datasets from the UK Biobank, including image-derived phenotypes (IDPs); focusing on white matter hyperintensities (WMHs) as an example of non-Gaussianity and a diffusion tensor imaging (DTI) modality for a whole-brain model.

Our proposed method makes it possible to apply normative modelling to considerably larger samples than was feasible before [7], [8]. The results from the computational experiments on the whole brain model showed that the BLR method is scalable to population-sized data sets and fine-grained voxel-level data. In comparison, most normative models used Gaussian process regression, which due to its high computational complexity could only be used in studies with a relatively low sample size. This improvement is mainly because the approximation of the covariance matrix by a set of basis functions allowed us to account for non-linearity in a computationally less demanding way than the Gaussian process regression method, therefore making the B-spline BLR scalable for big datasets. Computationally scalable modelling of nonlinear effects is important since our experiments showed that a cubic B-spline transformation of the age covariate improved model fit compared to linear models for most neuroimaging modalities.

Another major benefit of our method is the possibility of modelling non-Gaussian distribution by the use of the likelihood warping technique. This is important in general, as the aim of normative modelling is to accurately model the centiles of variation in addition to modelling the mean and is especially important for normative modelling of variables that are not approximately Gaussian distributed. For example, we showed that the WMHs show non-Gaussian behaviour that is well suited to uncover the benefits of the warped model over the standard model. We demonstrated the improved fit of the WMHs by including a B-spline transformation and a SinhArcsinh likelihood warping in the normative model, which was also exemplified for the longitudinal data. The same improvement in fit for other data modalities that showed more non-Gaussianity in their residuals was also demonstrated by comparing the warped BLR to the B-spline BLR for all the IDPs. Furthermore, it was shown on a whole-brain model of a DTI modality that for several voxels the warped BLR gives a better model performance than a B-spline BLR.

We emphasize that the addition of non-linear effects and non-gaussianity makes the model more flexible which increase the need for model selection in order to avoid possible overfitting. We presented several model selection criteria that can be used to choose the optimal model settings for different neuroimaging modalities. It should be recognized that for some IDPs and voxels the B-spline BLR gives a better fit, showing that a more flexible model is not always needed. Therefore, we recommend carefully examining the type of data one wants to model and based on the data trends found for the residuals (Gaussian or non-Gaussian) to decide if a more flexible model is preferred. This can easily be checked by looking at the skewness and kurtosis of the distribution or making a QQ-plot. Additionally, different model selection criteria can sometimes contradict each other, as they are sensitive to different parts of the data. As we showed above, classical metrics such as EV and MSLL are not very sensitive to the shape of the predictive distribution. The consequence is that per task, we have to decide if we want a better EV, most sensitive to the mean fit and dependent on the flexibility of the model, or a better MSLL/BIC, which is more sensitive to the variance and penalizes the flexibility of the model. The variability in model selection criteria demonstrates that for different imaging modalities, different normative modelling settings are preferred and the added flexibility is confirmed to only give an advantage for response variables that show non-Gaussianity in their residuals.

We confirmed that the deviations from the normative modelling frame-work can be meaningfully related to behaviour. We established a significant correlation between the warped deviance scores from the IDPs and several dimensions of the intelligence phenotype. These tests give a first indication of the possible relationships between the deviations and behaviour. For the whole brain model, the relationship with behaviour was shown with a significant correlation between an approximation to the g-factor in the form of the first principal component of the cognitive phenotypes and the warped deviance scores. This study demonstrates that the model could be extended to make predictive scores not only in the brain domain, but also for the behavioural phenotype. In the future, the neurobiological markers of deviation from normal development can be extended to become markers of psychiatric disorders. This has already been done on a smaller scale, using normative modelling [9], [10], [13], [30], [33], [34], but we would like to extend these studies to bigger data models, which include a wide variety of neuroimaging data modalities.

In conclusion, the current study suggests that non-linearity and non-Gaussianity are two parameters of big neuroimaging datasets that need to be captured to make accurate predictions for normal brain development. In this paper, we have done that through a warped BLR normative model. We have shown using several neuroimaging modalities the benefit of this model over more conservative models. Caution is essential when applying non-Gaussian models, as they can overfit and should mainly be used in the presence of non-normally distributed residuals. We recommend carefully assessing the distribution of residuals and the model selection parameters using the different model selection criteria mentioned in this paper that give a balance between model complexity and model fit.

## Appendix A

Figure A.8 shows the Bland-Altman plots of the explained variance for the IDPs and FreeSurfer measurements comparing the different model settings.

## Appendix B

An example list of the IDPs, processed using FUNPACK (the FMRIB UKBiobank Normalisation, Parsing And Cleaning Kit), used in this study is given in B.1. The IDPs contained the following neuroimaging modalities [17]:

1. T1, from which the total brain volumes are calculated.
2. Resting-state fMRI, from which the apparent connectivity between certain brain regions is estimated.
3. Task fMRI, from which the strength of response to certain tasks is given, which can be related to higher cognitive functioning.
4. T2 Flair, from which the white matter lesions are estimated.
5. DMRI, from which the DTI measures such as FA and MD are calculated.
6. Susceptibility-weighted imaging (SWI), from which venous vasculature, microbleed and other aspects of microstructure are estimated.

**Table B.1:**
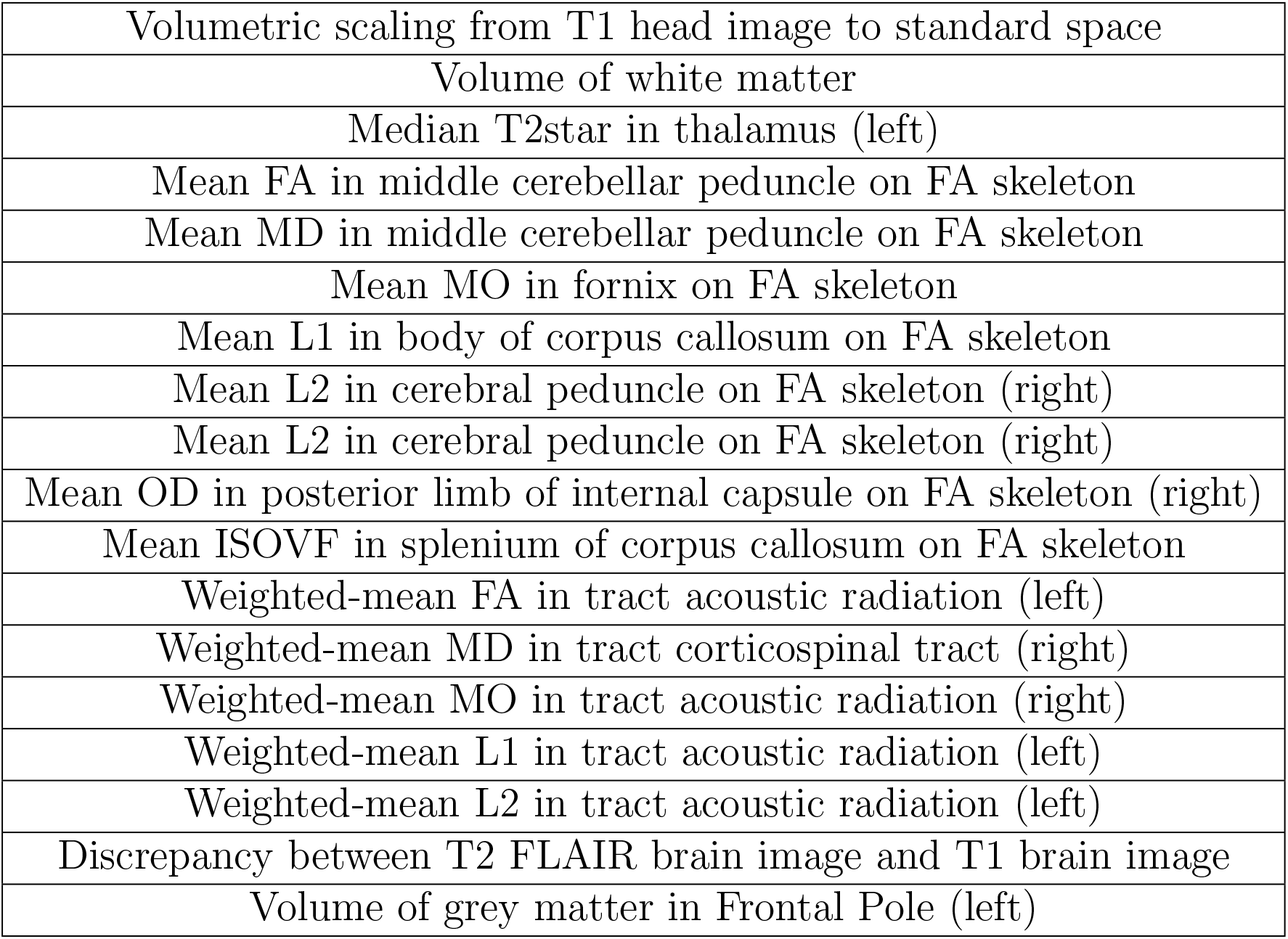
Example list of the IDP field names, processed using FUNPACK (the FMRIB UKBiobank Normalisation, Parsing And Cleaning Kit).

|  |
| --- |
| Volumetric scaling from T1 head image to standard space |
| Volume of white matter |
| Median T2star in thalamus (left) |
| Mean FA in middle cerebellar peduncle on FA skeleton |
| Mean MD in middle cerebellar peduncle on FA skeleton |
| Mean MO in fornix on FA skeleton |
| Mean L1 in body of corpus callosum on FA skeleton |
| Mean L2 in cerebral peduncle on FA skeleton (right) |
| Mean L2 in cerebral peduncle on FA skeleton (right) |
| Mean OD in posterior limb of internal capsule on FA skeleton (right) |
| Mean ISOVF in splenium of corpus callosum on FA skeleton |
| Weighted-mean FA in tract acoustic radiation (left) |
| Weighted-mean MD in tract corticospinal tract (right) |
| Weighted-mean MO in tract acoustic radiation (right) |
| Weighted-mean L1 in tract acoustic radiation (left) |
| Weighted-mean L2 in tract acoustic radiation (left) |
| Discrepancy between T2 FLAIR brain image and T1 brain image |
| Volume of grey matter in Frontal Pole (left) |

## Appendix C

We computed the differences between the BICs of a B-spline BLR and a warped BLR. Afterwards, we selected the top 30 IDPs where the B-spline model had the lowest BIC comparatively to the warped score or the other way around. In table C.2 the model selection criteria of the top 30 best-fitted IDPs with the B-spline BLR compared to the warped BLR are shown. In table C.3 the model selection criteria of the top 30 best-fitted IDPs with the warped BLR compared to the B-spline BLR shown. These tables demonstrate that every neuroimaging modality has its optimal model settings and that one should carefully examine the model selection criteria and shape of the response distribution, before choosing a model.

## Appendix D

We used a paired-sample t-test, pairing the IDP results (EV, MSLL and BIC) of the different models to estimate the difference between performance measures of the warped and non-warped BLR. In table D.4 and D.5 the Cohen’s d effect sizes and p-values are reported. The results show that there is a large difference between the standard BLR and the B-spline BLR, which confirms that one should take into account the non-linearity of the data. For the warped BLR and the B-spline BLR model, there is only a significant difference in the BIC score. We argue that this is because the model selection criteria are not necessarily sensitive to the deviations in the residuals from normality. Therefore, we also recommend to, alongside the model selection criteria, look at the skewness and kurtosis values together with the QQ-plot to choose the optimal model settings for each modality.

## Appendix E

In table E.6 we listed the cognitive variables from the UK Biobank that were used in this study with their IDs.

---

**Table C.2:**
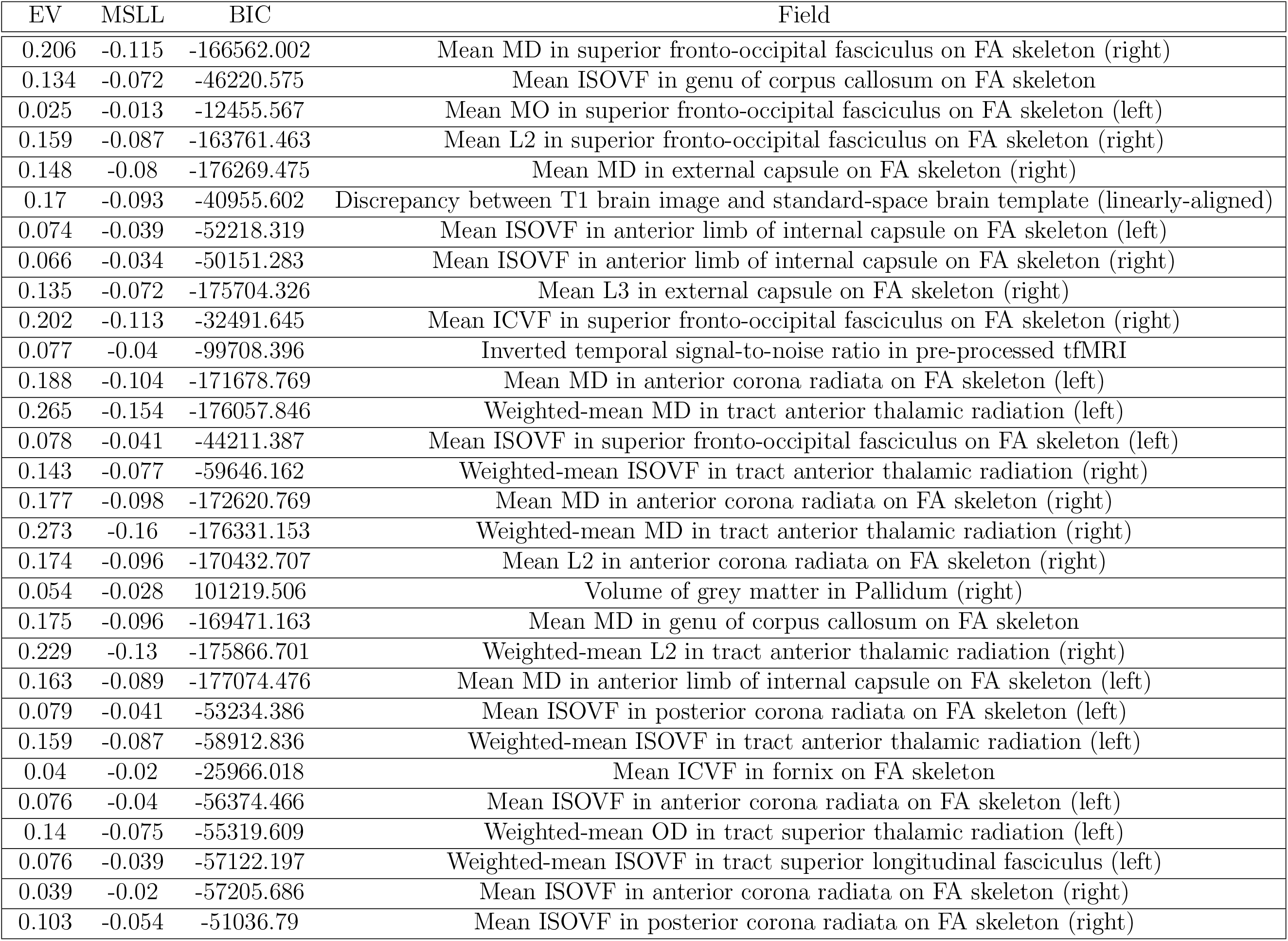
Model selection criteria of the top 30 IDPs, ranked according to difference between the BIC of a B-spline BLR and a SinhArcsinh warped BLR, where the B-spline BLR had a lower BIC score.

**Table C.3:** Model selection criteria of the top 30 IDPs, ranked according to the difference between the BIC of a B-spline BLR and a SinhArcsinh warped BLR, where the SinArcsinh warped BLR had a lower BIC score.

| EV | MSLL | BIC | Field |
| --- | --- | --- | --- |
| 0.249 | -0.143 | 184900.524 | Total volume of white matter hyperintensities (from T1 and T2-FLAIR images) |
| 0.147 | -0.079 | -29710.013 | Mean OD in fornix on FA skeleton |
| 0.285 | -0.164 | -137192.133 | Mean MD in fornix on FA skeleton |
| 0.276 | -0.153 | -136161.29 | Mean L3 in fornix on FA skeleton |
| 0.275 | -0.151 | -134595.545 | Mean L2 in fornix on FA skeleton |
| 0.153 | -0.083 | -87376.141 | Inverted temporal signal-to-noise ratio in pre-processed rfMRI |
| 0.27 | -0.157 | -24636.152 | Mean FA in fornix on FA skeleton |
| 0.171 | -0.093 | -32985.173 | Mean MO in anterior limb of internal capsule on FA skeleton (right) |
| 0.094 | -0.049 | -22330.216 | Mean MO in tapetum on FA skeleton (left) |
| 0.043 | -0.022 | -26681.768 | Mean MO in tapetum on FA skeleton (right) |
| 0.141 | -0.076 | -33305.028 | Mean MO in anterior limb of internal capsule on FA skeleton (left) |
| 0.054 | -0.027 | -42459.737 | Weighted-mean ISOVF in tract parahippocampal part of cingulum (left) |
| 0.117 | -0.062 | -71451.215 | Mean OD in splenium of corpus callosum on FA skeleton |
| 0.064 | -0.033 | -40476.534 | Weighted-mean FA in tract parahippocampal part of cingulum (right) |
| 0.307 | -0.183 | -15506.712 | Mean ISOVF in fornix on FA skeleton |
| 0.182 | -0.1 | -34039.973 | Discrepancy between T2 FLAIR brain image and T1 brain image |
| 0.047 | -0.024 | -41660.315 | Weighted-mean FA in tract parahippocampal part of cingulum (left) |
| 0.058 | -0.03 | -51125.932 | Mean OD in tapetum on FA skeleton (left) |
| 0.199 | -0.111 | -172072.977 | Weighted-mean MD in tract posterior thalamic radiation (left) |
| 0.311 | -0.186 | -26746.982 | Discrepancy between tfMRI brain image and T1 brain image |
| 0.131 | -0.071 | -169248.259 | Mean MD in posterior thalamic radiation on FA skeleton (left) |
| 0.089 | -0.046 | -181090.417 | Mean MD in inferior cerebellar peduncle on FA skeleton (left) |
| 0.07 | -0.036 | -41654.584 | Weighted-mean ISOVF in tract parahippocampal part of cingulum (right) |
| 0.028 | -0.014 | -35788.551 | Mean MO in posterior limb of internal capsule on FA skeleton (right) |
| 0.069 | -0.036 | -62423.772 | Weighted-mean OD in tract forceps major |
| 0.027 | -0.014 | -52538.461 | Mean ISOVF in middle cerebellar peduncle on FA skeleton |
| 0.314 | -0.188 | -27837.003 | Discrepancy between rfMRI brain image and T1 brain image |
| 0.085 | -0.044 | -170720.346 | Weighted-mean MD in tract medial lemniscus (right) |

**Table D.4:**
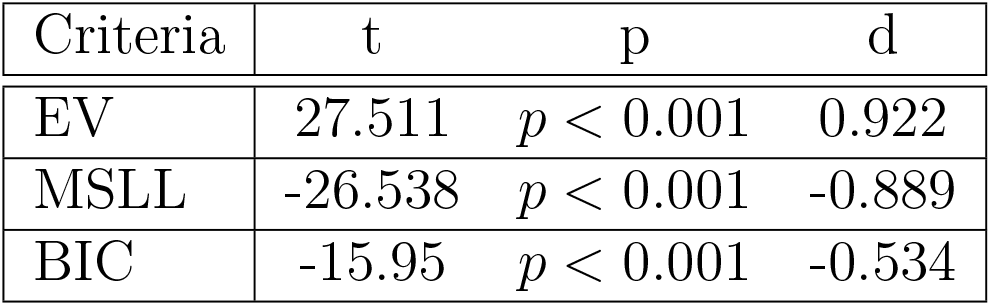
Table presenting a paired-sample t-test between the B-spline and standard BLR models, using the IDP data, showing a significant difference between the model selection criteria of the B-spline BLR and the standard BLR, with a large effect size.

| Criteria | t | p | d |
| --- | --- | --- | --- |
| EV | 27.511 | $p < 0.001$ | 0.922 |
| MSLL | -26.538 | $p < 0.001$ | -0.889 |
| BIC | -15.95 | $p < 0.001$ | -0.534 |

**Table D.5:**
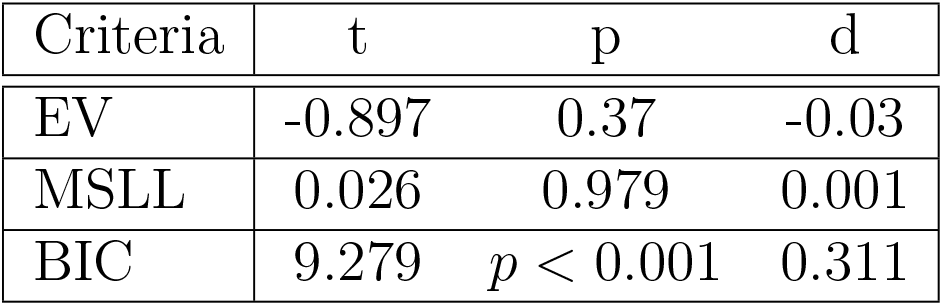
Table presenting a paired-sample t-test between the B-spline and warped BLR models, using the IDP data, showing only a significant difference between the model selection criteria of the B-spline BLR and the B-spline SinhArcsinh warped BLR using the BIC score, with a small effect size.

**Table E.6:** Cognitive variables of the UK Biobank that were used in this study.

| Field | FieldID |
| --- | --- |
| Number of times snap-button pressed | 403 |
| Duration to first press of snap-button in each round | 404 |
| Mean time to correctly identify matches | 20023 |
| Time elapsed | 4256 |
| Digits entered correctly | 4259 |
| Number of rounds of numeric memory test performed | 4283 |
| Time to complete test | 4285 |
| Duration screen displayed | 4290 |
| Number of attempts | 4291 |
| Prospective memory result | 20018 |
| Fluid intelligence score | 20016 |
| Number of fluid intelligence questions attempted within time limit | 20128 |
| Duration to complete numeric path (trail 1) | 6348 |
| Total errors traversing numeric path (trail 1) | 6349 |
| Duration to complete alphanumeric path (trail 2) | 6350 |
| Total errors traversing alphanumeric path (trail 2) | 6351 |
| Errors before selecting correct item in numeric path (trail 1) | 6770 |
| Errors before selecting correct item in alphanumeric path (trail 2) | 6771 |
| Interval between previous point and current one in numeric path (trail 1) | 6772 |
| Interval between previous point and current one in alphanumeric path (trail 2) | 6773 |
| Number of puzzles correctly solved | 6373 |
| Number of puzzles viewed | 6374 |
| Number of puzzles correct | 6382 |
| Number of puzzles attempted | 6383 |
| Number of puzzles correct | 21004 |
| Number of symbol digit matches attempted | 23323 |
| Number of symbol digit matches made correctly | 23324 |

